# Contributions of spatial and temporal control of step length symmetry in the transfer of locomotor adaptation from a motorized to a non-motorized split-belt treadmill

**DOI:** 10.1101/699736

**Authors:** Daniel L. Gregory, Frank C. Sup, Julia T. Choi

## Abstract

**Background:** Locomotor adaptation during motorized split-belt walking depends on independent processes for spatial and temporal control of step length symmetry. The unique mechanics of motorized split-belt walking that constrains two limbs to move at different speeds during double support may limit transfer of step length adaptations to new walking contexts.

**Research question:** How do spatial and temporal locomotor outputs contribute to transfer of step length adaptation from constrained motorized split-belt walking to unconstrained non-motorized split-belt walking?

**Methods:** We built a non-motorized split-belt treadmill that allows the user to walk at their own pace while simultaneously allowing the two belts to be self-propelled at different speeds. 10 healthy young participants walked on the non-motorized split-belt treadmill after an initial 10-minute adaptation on the motorized split-belt with a 2:1 speed ratio. Foot placement relative to the body and timing between heel strikes were calculated to determine spatial and temporal motor outputs, respectively. Separate repeated measures ANOVAs were used for step length difference and its spatial and temporal components to assess for transfer to the non-motorized treadmill.

**Results:** We found robust after-effects in step length difference during transfer to non-motorized split-belt treadmill walking that were primarily driven by changes in temporal motor outputs. Conversely, residual after-effects observed during motorized tied-belt treadmill walking (post-transfer) were driven by changes in spatial motor outputs.

**Significance:** Our data showed decoupling of adapted spatial and temporal locomotor outputs during the transfer to non-motorized split-belt walking, raising the new possibility of using a non-motorized split-belt treadmill to target specific spatial or temporal gait deficits.

## Introduction

Control of human locomotion is highly adaptable. Gait patterns can be altered through motorized treadmill training, even after central nervous system damage [1]. Walking on a split-belt treadmill induces asymmetric step lengths by constraining limbs to move at different speeds which alters gait through an error-based learning process known as motor adaptation [2], [3]. Adaptation of symmetric step lengths during motorized split-belt treadmill walking results from changes in both spatial control of foot placement, and temporal control of step timing [4]–[6]. These spatial and temporal motor outputs reflect distinct neural strategies to overcome the asymmetric belt speeds. Positive asymmetries in foot placement and step timing achieved during adaptation persist during de-adaptation, while the distance the foot travels during stance immediately returns to tied baseline values [6]. This after-effect during post-adaptation walking reflects the updating of neural control processes for walking and is limited when transferring to over-ground walking or to different modes of locomotion [7]–[11].

Step length asymmetry post-stroke is associated with both spatial and temporal abnormalities in gait control [3], [4], [12], [13]. Training on a motorized split-belt treadmill is a promising rehabilitation strategy to reduce asymmetric step length patterns in hemiparetic walkers [3], [14]. After repeated exposure to split-belt adaptation, spatial asymmetries to walking control improve substantially and persist up to three months after an intervention. However, temporal asymmetries such as stance time and double support time remain unchanged both immediately and after 3 months follow up [15]. Further, changing the context of the walking condition substantially reduces or eliminates transfer in both post-stroke [7], and healthy individuals [8]. Potentially limiting transfer of motorized split-belt treadmill adaptations to over-ground walking is the unique mechanics of split-belt treadmills—the constraint to have limbs move at different speeds during the double support period.

To address these issues and better understand the capacity to transfer locomotor adaptations, we developed a non-motorized split-belt treadmill that allows the user to determine their own walking pace while simultaneously allowing for asymmetric behavior. In contrast to motorized treadmills, non-motorized treadmills have freely moveable belts which are driven by participants pushing against an inclined or concave surface which allow participants to self-select and express natural gait variability [16]–[18]. Our custom non-motorized split-belt treadmill is also cheaper, more portable, and safer compared to motorized systems. Prior research on non-motorized treadmills for locomotor adaptation is limited to single-belt devices [19]. A better understanding of the generalization between motorized and non-motorized split-belt devices would allow us to implement split-belt therapy using lower-cost non-motorized systems, and thereby improve the translational potential of treadmill training to community ambulation.

The current study is first to examine (1) whether step length adaptation from motorized split-belt treadmill walking transfers to non-motorized treadmill walking, and (2) how the separate spatial and temporal control of step length symmetry contributes to transfer in a novel walking context. Non-motorized treadmills offer a unique combination of portability and economy which we believe will provide a valuable bridge between split-belt treadmill and over ground walking. We hypothesized that after-effects would be present during transfer and would be driven by positive asymmetries in foot placement and step timing. We further hypothesized that after-effects washed out during non-motorized treadmill walking should lead to diminished or absent after-effects during tied motorized treadmill walking.

## Methods

### Participants

Ten healthy volunteers (six female, four male; Age 26.5 ± 5.6 years) with no neurological or biomechanical impairments were recruited for this study. All study protocols were approved by the University of Massachusetts, Amherst Institutional Review Board. All participants provided written informed consent prior to enrollment. None of the participants had prior experience walking on a split-belt treadmill.

### Experimental setup

#### Motorized split-belt treadmill

Participants walked on a split-belt treadmill (Bertec Corp, Columbus OH) that has separate left and right belts, each with its own motor. During motorized treadmill walking, participants wore a non-weight-bearing safety harness suspended from the ceiling. Participants were instructed to minimize handrail use, walking with normal arm swing, and to maintain forward gaze.

#### Non-motorized spit-belt treadmill

We designed and built a user-propelled non-motorized split-belt treadmill (fig. 1a). The non-motorized treadmill was fabricated from two commercially available non-motorized treadmills (Inmotion II Manual Treadmill; Stamina, Springfield, MO), which were designed to share a common support structure and minimize spacing between the belts (32 mm). To minimize friction with the belt-deck interface, a sheet of polytetrafluoroethylene (PTFE, 0.30” sheet; ePlastics, San Diego, CA) plastic was secured each treadmill deck. A ~1kg mass was added to each flywheel to increase the inertial properties of the belt-flywheel system to ensure continuous movement as the limb transitioned from stance to swing, enabling smoother transition into the next stance phase. The treadmill utilizes gravity (~13° incline) and the users body weight to assist driving the symmetrically resisted but independently user-propelled belts. During non-motorized treadmill walking, participants lightly held the handrails and were instructed to avoid supporting body weight.

**Figure 1.**
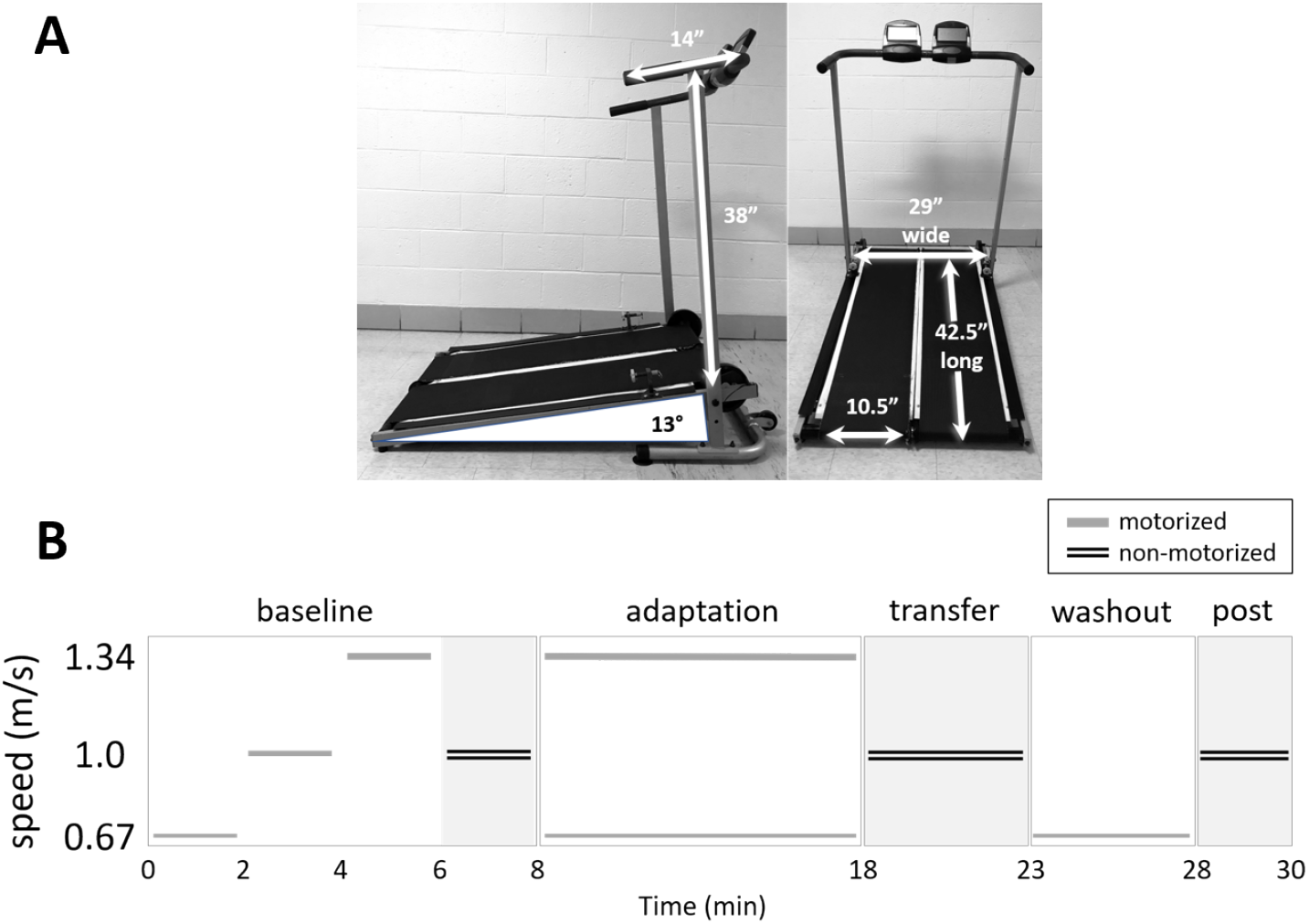
Non-motorized split-belt treadmill and paradigm. ***A.*** Non-motorized treadmill dimensions (inches), *left panel*; handrail height from top of treadmill deck and handle depth, and deck angle in degrees. Non-motorized treadmill dimensions, *right panel*; deck width and length and individual belt widths. ***B.*** Split-belt treadmill trasnfer paradigm. All subjects were recorded in baseline conditions for slow, medium, fast, and non-motorized treadmill (familiarization, not recorded and not shown) conditions in random order. The split-belt adaptation period was always immediately preceded by a second non-motorized treadmill baseline period used for non-motorized treadmill baseline measures. After all baseline periods were completed, all subjects adapted for 10 minutes with one limb on a belt traveling at the slow speed (0.67 m/s) and the other at the fast speed (1.34 m/s). Subjects then moved to the non-motorized treadmill without forward stepping to assess for the transfer to the non-motorized treadmill. Transfer was 5 minutes at preferred speed. Subjects then moved without forward stepping back to the motorized treadmill to assess for residual after-effects specific to the motorized treadmill for an additional 5 minutes. Finally, subjects then returned to the non-motorized once more to ensure all residual after-effects had been washout out for an additional 2 minutes.

### Split-belt adaptation paradigm

The adaptation paradigm (fig. 1b) consisted of motorized tied-belt, motorized split-belt and non-motorized split-belt walking conditions. During Baseline, participants walked on the motorized treadmill with belts tied at a slow (0.67 m/s), medium (1.0 m/s), and fast (1.34 m/s) speeds for two minutes each, randomized across participants. A 5-minute non-motorized treadmill Baseline trial was collected at preferred walking speed. Participants were instructed to “walk as fast as you would to a meeting for which you have adequate time to arrive”. During *Adaptation*, participants walked on the motorized split-belt at a 2:1 speed ratio (0.67 and 1.34 m/s) for ten minutes. These speeds were chosen because the average speed is 1.0 m/s, which was the average preferred walking speed on the non-motorized treadmill during pilot testing. Participants were then instructed to side-step from the motorized treadmill onto the non-motorized treadmill less than 2-feet away. Forward stepping was avoided to prevent washout of after-effects. During *Transfer*, participants walked for 5 minutes on the non-motorized treadmill. Participants returned to the motorized treadmill for the *Washout* period and walked at the slow speed for an additional 5 minutes. During *Post-adaptation*, participants again walked on the non-motorized treadmill for 2 minutes. Participants were randomly assigned left or right limb to the slow belt during split-belt walking and subsequent references to a limb will be as slow or fast regardless of condition.

### Data collection

During motorized treadmill walking, kinematics were recorded at 100 Hz using a 4-camera motion capture system (Qualisys, Sweden). Reflective markers were placed bilaterally over the fifth metatarsal, lateral malleolus, tibial plateau, greater trochanter, and the anterior superior iliac spines. During non-motorized treadmill walking, kinematics were relegated to use of only ankle markers for identification of spatio-temporal parameters due to limitations in capture space during data collection.

### Data analysis

Data analysis was performed in MATLAB (Mathworks, Natick, MA, v. R2017a). Marker data were low pass filtered at 6 Hz with a second order Butterworth filter. For both motorized and non-motorized treadmill walking, we defined heel contact and toe-off as the time of peak anterior and posterior ankle position for each step, respectively. We used step length difference to assess split-belt walking adaptation [2], [3], [14], [20], using the anterior-posterior distance between the ankle markers at the time of heel contact. Step length difference was defined as SL_fast_ – SL_slow_, where SL_fast_ is the step length with fast leg leading and SL_slow_ is with the slow leg leading at heel strike (fig. *2a*). We also included double support difference as a secondary outcome (see Supplementary materials).

**Figure 2.**
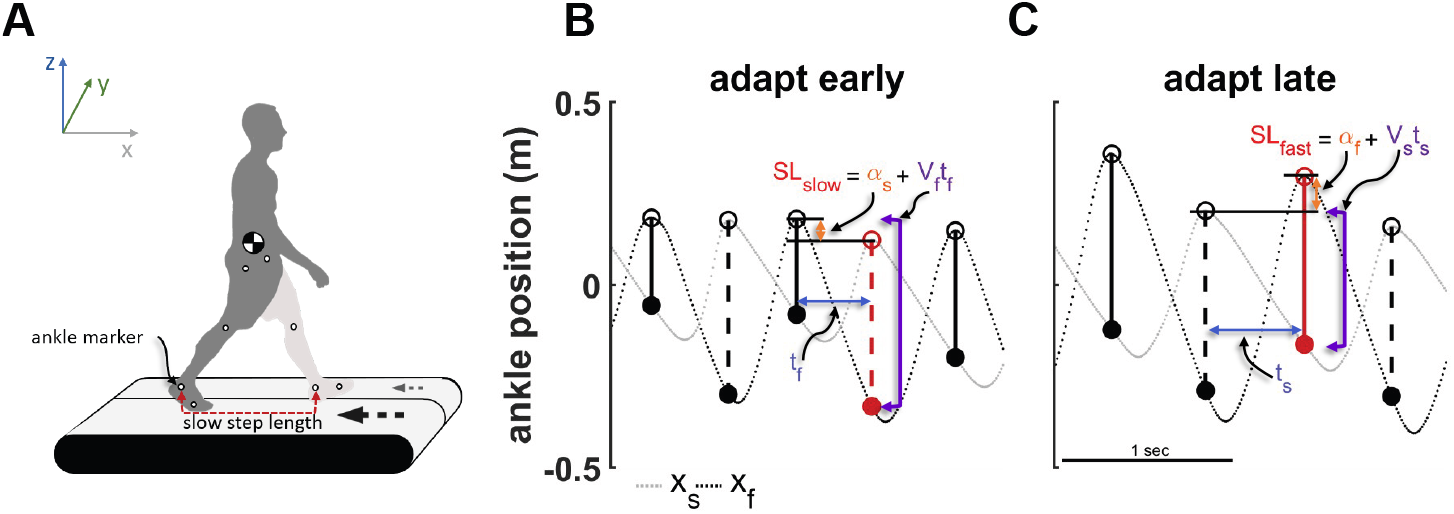
Analytical calculation of step length. Visual representation of walking parameters for the analytical step length derivation. ***A.*** Walker demonstrating the position of the reflective markers on the fifth metatarsal, lateral malleolus, and lateral condyle. Gray crossed circle represents the pelvic centroid used to calculate the reference frame for ankle marker data on the motorized treadmill. Slow instantaneous step length shown (red arrow line) as distance between ankle markers at time of slow limb heel strike. Gray and black arrows indicate belt speeds. ***B.*** Slow step length visualized during early adaptation. Ankle marker trajectories in the pelvic centroid reference frame. Positive values indicate positions in front of the pelvic centroid and negative indicate positions behind the pelvic centroid. Dashed lines indicate individual ankle marker trajectories for the fast (black, x_f_), and slow (gray, x_s_) limbs. Ankle marker trajectories are plotted with time on the x-axis, therefore distance between dots indciates distance traveled every 10 milliseconds. Open circles indicate the position and time of the respective limbs heel strike. Filled circles indicate the position and time of the trailing limb ankle at the leading limbs heel strike. Solid vertical lines connecting open and closed circles indicate the instantaneous fast step length and the vertical dashed lines connecting open and closed circles indicate the instantaneous slow step length (i.e. SL_slow_). The slow analytical step length (dashed red line) is the summation of the (negative) spatial component (α_s_), and the product of the velocity of the fast limb (v_f_) and the fast step time (t_f_), the distance traveled by the fast limb from fast heel strike to fast position at slow heel strike. v_f_ is the difference between the fast limb ankle (x_f_) at fast heel strike, and the fast limb ankle at slow limb heel strike, divided by the fast step time (t_f_). ***C.*** Fast step length visualized dyring late adaptation. The fast analytical step length (solid red line) is the summation of the spatial component (α_f_), and the product of the velocity of the slow limb (v_s_) and the slow step time (t_s_).

To quantify the spatial and temporal contribution to changes in step length, we applied the following analytical model of step length difference (for derivation, see Finley et al. 2015):

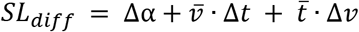

where the first term (step position) represents the difference between the relative positions of the feet at the fast and slow heel strikes, the second term (step time) is the difference in slow and fast step times as a function of average foot speed, and the third term (step velocity) is the difference in slow and fast foot velocities as a function of the average step time (fig. *2b,c*). It is important to note that during motorized treadmill conditions, fast and slow speeds are fixed, whereas during non-motorized treadmill walking, the speed of each belt/limb is strictly user controlled and becomes a third degree of freedom while walking.

### Statistical analysis

Group means were calculated in bins across the last 10 strides for each Baseline trial (motorized slow, motorized medium, motorized fast, and non-motorized preferred) and the first 5 strides (early) and last 5 strides (late) for Adaptation, Transfer, Washout, and Post trials. Separate repeated measures ANOVAs were used to test the effects of time on each dependent variable. Post-hoc analyses were performed using independent sample t-tests to compare each dependent variable at each time point to the baseline value and for comparisons between early and late time points.

## Results

### Adaptation

Repeated measures ANOVA revealed significant effects of time on step length (F(7,63) = 25.17, p < 0.001, fig. 3a–5a). Group averaged step-length differences were not significantly different from zero during baseline tied-walking on the motorized treadmill (all p-values > 0.05). Figure 3a shows gradual changes in step length difference (dark grey/black) during the first 100 and last 20 strides in Adaptation. During *early Adaptation*, step length difference showed a large initial change compared to baseline (p < 0.001, fig. 3b). By *late Adaptation*, step length difference was reduced back to baseline levels (p = 0.58, fig. 3b).

**Figure 3.**
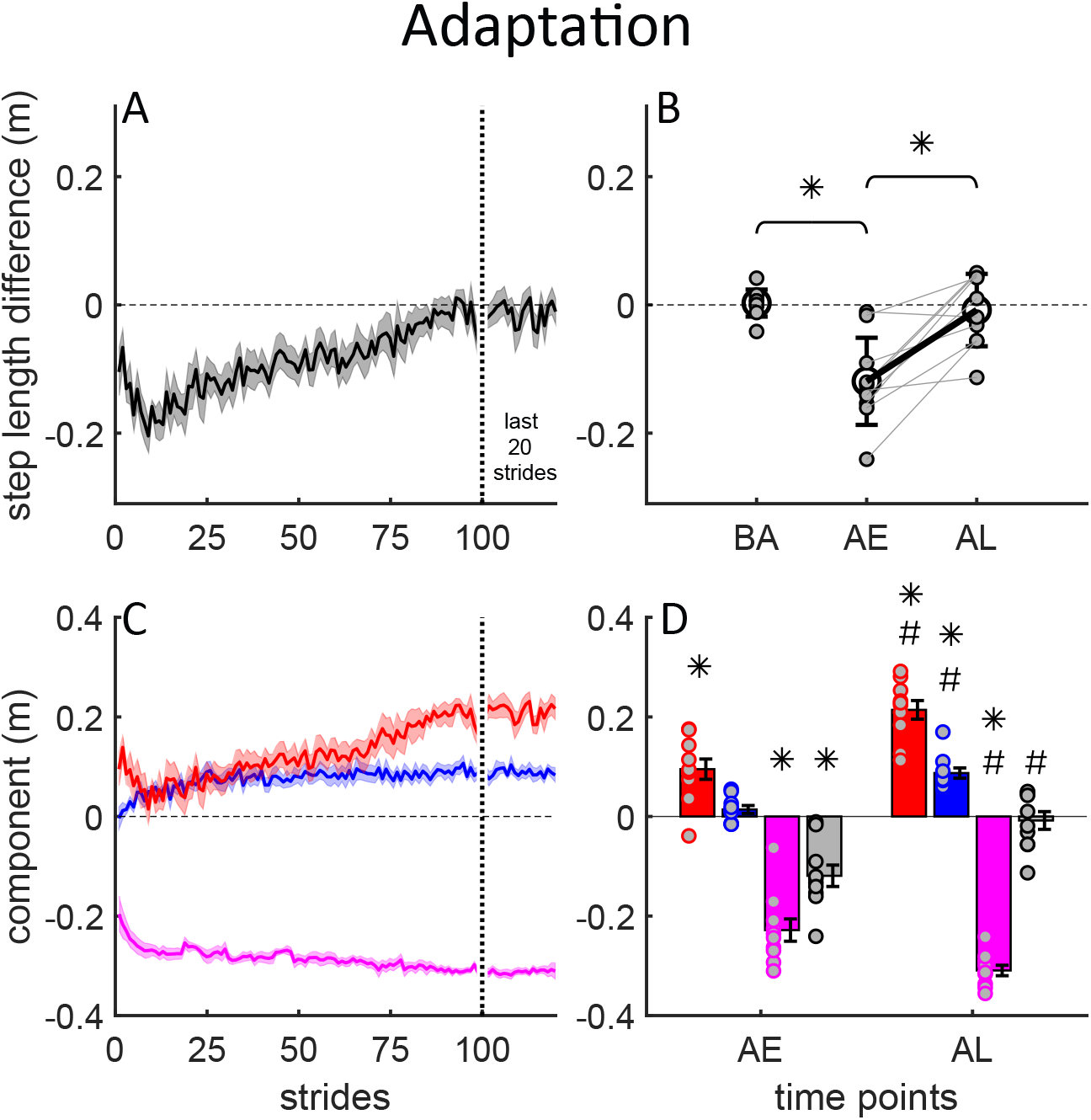
Adaptation; Progression of step length difference and contribution of spatial, temporal, and velocity asymmetries to motorized split-belt treadmill walking during adaptation. ***A.*** Group average step length difference during the first 100 and last 20 strides of the adaptation period. Shaded area represents plus/minus standard error. Negative values indicated the slow step length is larger than the fast step length. ***B.*** Individual subject step length difference data (small gray filled circles) comparing baseline (BA) slow walking to early adapt (AE) and late adapt (AL). Light gray lines connect individual subject data points between AE and AL. Group means are represented by large black circles, error bars are standard deviations. Thick black line connecting AE and AL indicates group change. Horizontal bars with star indicate significant differences between time points (p < 0.05). ***C.*** Stride-by-stride changes in individual components for the first 100 and last 20 strides; spatial (red), temporal (blue), and velocity (magenta) differences. Shaded areas are plus/minus standard error. ***D.*** Individual subject component data (gray filled colored circles) comparing AE and AL. Bars are group means, error bars are standard error. “*” indicates significant difference to baseline slow, “#” indicates late time point significantly different from early (p < 0.05).

As previously reported [4]–[6], step length difference can be decomposed into independent step velocity (magenta), step position (red) and step time (blue) contributions (fig 3c,d). The step velocity component became increasingly negative over the course of adaptation, reflecting a large negative velocity induced perturbation which required opposition (p < 0.001, fig 3d). During *early Adaptation*, the step position component showed a significant initial asymmetry (p < 0.001, fig. 3d). The step time component showed a more gradual change, with *early Adaptation* values not significantly different from baseline (p = 0.26, fig. 3d). By *late Adaptation*, the step position and step time components both increased significantly (p < 0.001, fig. 3d) to cancel the negative step velocity component, resulting in symmetrical step lengths (p = 0.58, fig. 3d).

### Transfer

We found significant transfer of motorized treadmill adaptations to the non-motorized split-belt treadmill. Figure 4a shows the after-effect in step length difference during the first 100 and last 20 strides in Transfer. During *early Transfer*, step length difference showed a negative after-effect relative to baseline (p < 0.001, fig. 4b). By *late Adaptation*, step length difference returned to baseline levels (p = 0.31).

**Figure 4.**
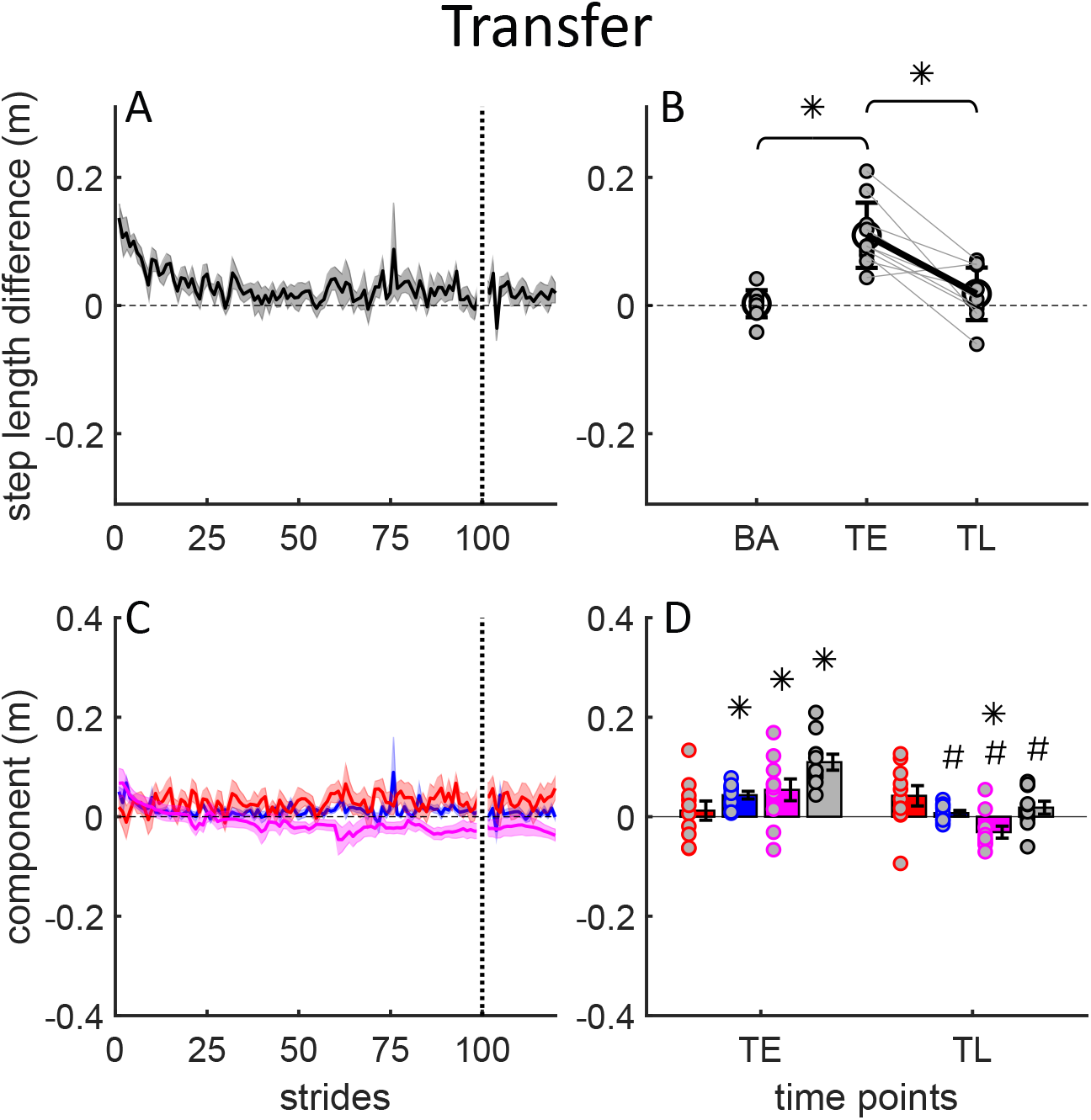
Transfer; Progression of step length difference and contribution of spatial, temporal, and velocity asymmetries to non-motorized split-belt treadmill walking during transfer. ***A.*** Group average step length difference during the first 100 and last 20 strides of the transfer period. Shaded area represents plus/minus standard error. Positive values indicated the fast step length is larger than the slow step length. ***B.*** Individual subject step length difference data (small gray filled circles) comparing baseline (BA) walking to early transfer (TE) and late transfer (TL). Light gray lines connect individual subject data points between TE and TL. Group means are represented by large black circles, error bars are standard deviations. Thick black line connecting TE and TL indicates group change. Horizontal bars with star indicate significant differences between time points (p < 0.05). ***C.*** Stride-by-stride changes in individual components for the first 100 and last 20 strides; spatial (red), temporal (blue), and velocity (magenta) differences. Shaded areas are plus/minus standard error. ***D.*** Individual subject component data (gray filled colored circles) comparing TE and TL. Bars are group means, error bars are standard error. “*” indicates significant difference to baseline non-motorized, “#” indicates late time point significantly different from early (p < 0.05).

Figure 4c,d show the contribution of the step velocity, position and step timing components to the step length after-effects during Transfer. In *early Transfer*, the step velocity component displayed a large positive (i.e. an after-effect) component that was significantly different from baseline (p = 0.03, fig. 4d, magenta). By *late Transfer*, the step velocity component decreased to a negative value that was significantly different from baseline (p = 0.02), indicating that participants settled on a step velocity difference in the same direction as the initial motorized split-belt perturbation (i.e. a negative velocity difference). The step position component was not significantly different from baseline during *early Transfer* (p = 0.5, fig. 4d). There were also no significant differences between *early* and *late Transfer* for the step position component (p = 0.3). By *late Adaptation*, the step position component was not different from baseline (p = 0.06, fig. 4d), however, upon inspection of individual data, this result may have been driven by a single outlier (*late Transfer*, red filled circles). These results indicate that the spatial component did not contribute to the initial after-effects in step length difference. The step time component was significantly different from baseline during *early Transfer* (p < 0.001, fig. 4d). By *late Transfer*, the step time component decreased back to baseline values (p = 0.58), indicating washout of the temporal after-effect.

### Washout

After-effects in step length reappeared during motorized treadmill walking in *early Washout* (p < 0.001). Step length difference was significantly different from baseline during early Washout (p < 0.001 fig. 5a) and returned back to baseline by *late Washout* indicated by significant differences from *early* to *late Washout* (p < 0.001) but did not completely return to baseline values (p = 0.03, fig. 5b).

**Figure 5.**
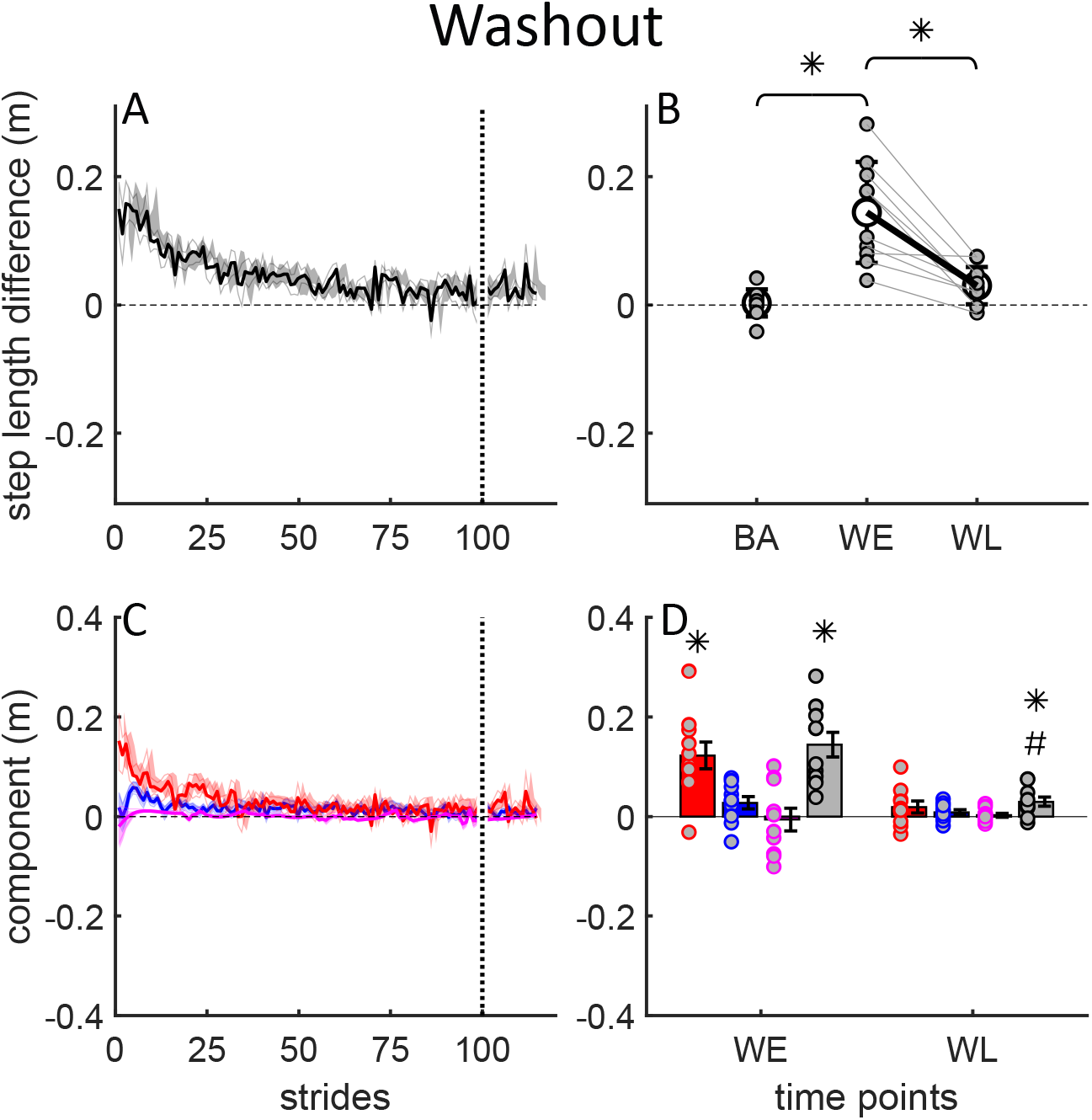
Washout; Progression of step length difference and contribution of spatial, temporal, and velocity asymmetries to motorized treadmill walking during washout. ***A.*** Group average step length difference during the first 100 and last 20 strides of the adaptation period. Shaded area represents plus/minus standard error. Positive values indicated the fast step length is larger than the slow step length. ***B.*** Individual subject step length difference data (small gray filled circles) comparing baseline (BA) slow walking to early washout (WE) and late washout (WL). Light gray lines connect individual subject data points between WE and WL. Group means are represented by large black circles, error bars are standard deviations. Thick black line connecting WE and WL indicates group change. Horizontal bars with star indicate significant differences between time points (p < 0.05). ***C.*** Stride-by-stride changes in individual components for the first 100 and last 20 strides; spatial (red), temporal (blue), and velocity (magenta) differences. Shaded areas are plus/minus standard error. ***D.*** Individual subject component data (gray filled colored circles) comparing WE and WL. Bars are group means, error bars are standard error. “*” indicates significant difference to baseline slow, “#” indicates late time point significantly different from early (p < 0.05).

Figure 5c shows the contribution of the step velocity, step position and step timing components to the step length after-effects during the Washout period. The step velocity components were not different from baseline during *early Washout* (p = 0.72) or *late Washout* (p = 0.95), due to the symmetric belt speeds (fig. 5c,d). The step position component displayed significant after-effects during *early Washout*, (p < 0.001) that reduced back to baseline by *late Washout* (p = 0.17, fig. 5d). The step time component was not significantly different from baseline during *early Washout* (p = 0.09) nor *late Washout* (p = 0.48, fig 5d, blue).

### Unlearning interaction

Our linear regression analysis showed that step length error magnitude during *early Transfer* did not predict the step length error magnitude during *early Washout* (p = 0.782, fig. 6), suggesting that transfer did not result in unlearning of motorized treadmill adaptation. Linear regression analysis on the individual components also showed no interaction between *early Transfer* and *early Washout* for the step position (p = 0.132), step time (p = 0.451), or velocity components (p = 0.457).

**Figure 6.**
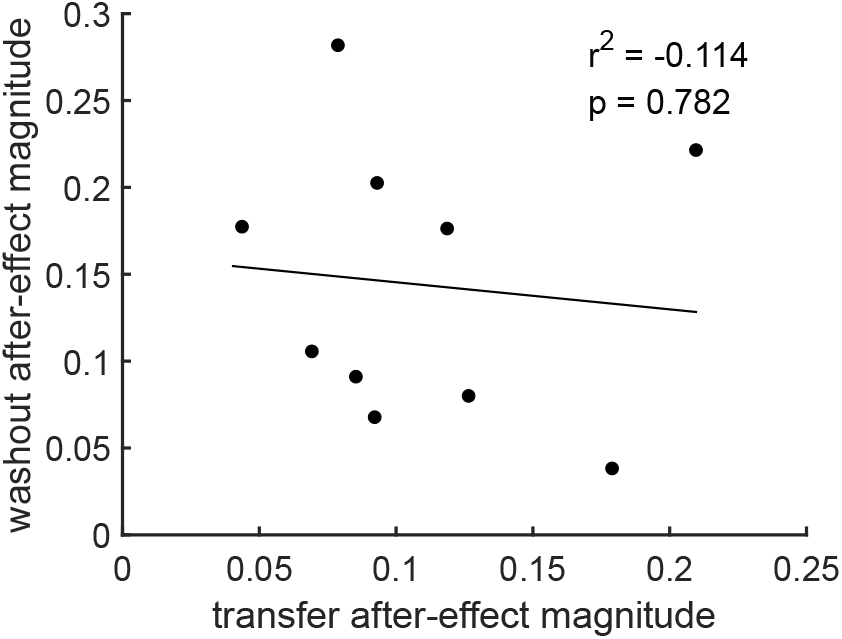
Linear regression to assess interaction between transfer after-effect and washout after-effect magnitude. Data points are individual subject averages from first 5 strides during early transfer (TE) and washout (WE), respectively.

### Post-washout

No residual step length difference after-effects were present once returned to the non-motorized treadmill post-washout (p > 0.05, not shown). In addition, there were no asymmetries between any individual component during *early Post-washout* walking (p > 0.05).

## Discussion

Effective movement rehabilitation requires the transfer of improvements made in the clinic to the community. Context is known to play a major role in modulating transfer of sensorimotor adaptations between different environments. Specifically, locomotor adaptation on a split-belt treadmill has repeatedly demonstrated limited transfer to over-ground walking in healthy and post-stroke participants [7], [8], [15], [21]. It is unclear based on previous studies whether spatial and temporal outputs contribute equally to transfer of split-belt locomotor adaptation. In agreement with our first hypothesis, we demonstrated transfer of motorized split-belt treadmill adaptation to non-motorized split-belt treadmill walking—a result driven by temporal and velocity but not spatial control asymmetries. These findings are promising and suggest that using a portable, low-cost split-belt treadmill for adaptive training can facilitate transfer from clinical to community walking.

Next, we hypothesized that the washout of after-effects during the Transfer period would lead to smaller or no after-effects during the Washout period [7]. However, we found large after-effects in both Transfer and Washout that were driven by asymmetries in step timing and foot placement differences, respectively. We suggest three potential mechanisms contributing to persistent after-effects. First, if the spatial and temporal parameters were uncoupled, this would suggest independent access to the control of spatial and temporal motor outputs and independent washout of each parameter. While previous work has demonstrated uncoupling of spatial and temporal control using feedback during adaptation [5], [6], the proposed uncoupling here occurred during the Transfer and Washout conditions. On the non-motorized treadmill, visual and tactile cues of the handrail and small walking surface may have led to an unintentional restriction of foot placement asymmetry, potentially blocking transfer of the spatial asymmetry. In line with this idea, there were no significant changes in foot placement differences during transfer as step length after-effects were a result of step time and belt velocity asymmetries. This suggests that by restricting spatial control during Transfer, temporal control was free to vary and likely washed out, leaving the spatial motor output to be deadapted during the Washout condition. Recent studies indicate that spatial and temporal control of gait may be related to functionally distinct neural circuitries. Spatial control has been associated with both state-estimation [22], a cerebellar process, and pre-planned cognitive control (i.e. cortical processes) in challenging walking environments [23]. In cerebellar lesion patients, double support difference, but not step length symmetry displays adaptive behavior [24]. Hemispherectomy patients adapt step lengths, but not double support difference in split-belt walking [14]. Together, these studies suggest that decoupling and subsequent independent washout of the temporal and spatial components of locomotion is possible if constraints limit access to the neural circuits which are involved in the control of a specific parameter. Our findings here are in line with this idea and suggest that uncoupling of spatial and temporal control is dissociable and may be implicitly and independently accessed.

Next, the apparent uncoupling of spatial and temporal after-effects could be a re-organization of spatial, temporal, and velocity asymmetries resulting from an additional degree of freedom (i.e. belt velocity) when walking on the non-motorized treadmill. This may effectively save the adapted pattern to be later washed out. Close inspection of the individual components during Transfer (Fig. 4c,d) might suggest a trade-off in spatial, temporal, and velocity components, which sum to symmetric step lengths. By late Transfer, belt speeds were asymmetric in the same direction as during Adaptation, while the spatial component had a positive asymmetry (Fig. 4d). Because the context and dynamics of each treadmill were similar, this would suggest that participants persisted with their adapted state from the motorized split-belt treadmill to trade-off symmetric step times and lengths with asymmetries in foot placement and step velocity. Consistent with this idea, previous work has demonstrated the high metabolic cost of locomotion with asymmetric step times [25]. Further, step length asymmetry alone during split-belt walking does not explain the added cost of asymmetry [26]. This may suggest that participants preferred temporal symmetry rather than the foot placement or velocity symmetry as a means of reducing metabolic cost. This further suggests that participants adapted a newly learned motor pattern to a novel condition in a way that was more efficient while meeting the demands of the novel environment, all while saving their newly adapted motor pattern for washout during the subsequent condition.

Last, partial washout or persistent after-effects during Transfer and Washout could also be caused by after-effects associated with different walking speeds between conditions [27]. The contribution of speed to after-effect size has been demonstrated previously in both over-ground and treadmill walking, with diminishing after-effects at both slower and faster speeds [10], [27]. In the current study, we show robust after-effects during both Transfer and Washout—independent of speed—with walking speed on the non-motorized treadmill being greater than the slow motorized speed (0.8 m/s versus 0.67 m/s, p < 0.05). Moreover, we demonstrated no association between Transfer and Washout after-effects, suggesting partial washout did not occur. Therefore, we reject the hypothesis that persistent after-effects were a function of different walking speeds.

In conclusion, we demonstrated that temporal (but not spatial) control contributed to the transfer of step length adaptation from motorized to non-motorized split-belt walking. One implication is that the non-motorized system could help translate split-belt training into standard of care because it has the advantage of being low cost, portable, and self-paced. A limitation of this study is that we do not have the data to directly compare after-effects in over-ground transfer. Further research is needed to investigate how non-motorized split-belt treadmills can be used in clinical populations, and whether training on this device would lead to improved over-ground walking.

## Supporting information

Supplemental material: Secondary outcome

## Acknowledgements

We thank Erin Carey, Jeffrey Florek, Nicholas Lake, Subhash Vallala, and Eric Murray for designing and building the non-motorized split-belt treadmill.

## References

[1] S. J. Mulroy, T. Klassen, J. K. Gronley, V. J. Eberly, D. A. Brown, and K. J. Sullivan, “Gait Parameters Associated With Responsiveness to Treadmill Training With Body-Weight Support After Stroke: An Exploratory Study,” Phys Ther, vol. 90, no. 2, pp. 209–223, Feb. 2010.

[2] D. S. Reisman, H. J. Block, and A. J. Bastian, “Interlimb Coordination During Locomotion: What Can be Adapted and Stored?,” Journal of Neurophysiology, vol. 94, no. 4, pp. 2403–2415, Oct. 2005.

[3] D. S. Reisman, R. Wityk, K. Silver, and A. J. Bastian, “Locomotor adaptation on a split-belt treadmill can improve walking symmetry post-stroke,” Brain, vol. 130, no. 7, pp. 1861–1872, Jul. 2007.

[4] J. M. Finley, A. Long, A. J. Bastian, and G. Torres-Oviedo, “Spatial and Temporal Control Contribute to Step Length Asymmetry During Split-Belt Adaptation and Hemiparetic Gait,” Neurorehabilitation and Neural Repair, vol. 29, no. 8, pp. 786–795, Sep. 2015.

[5] A. W. Long, R. T. Roemmich, and A. J. Bastian, “Blocking trial-by-trial error correction does not interfere with motor learning in human walking,” Journal of Neurophysiology, vol. 115, no. 5, pp. 2341–2348, Feb. 2016.

[6] L. A. Malone, A. J. Bastian, and G. Torres-Oviedo, “How does the motor system correct for errors in time and space during locomotor adaptation?,” Journal of Neurophysiology, vol. 108, no. 2, pp. 672–683, Apr. 2012.

[7] D. S. Reisman, R. Wityk, K. Silver, and A. J. Bastian, “Split-Belt Treadmill Adaptation Transfers to Overground Walking in Persons Poststroke,” Neurorehabil Neural Repair, vol. 23, no. 7, pp. 735–744, Sep. 2009.

[8] G. Torres-Oviedo and A. J. Bastian, “Seeing Is Believing: Effects of Visual Contextual Cues on Learning and Transfer of Locomotor Adaptation,” Journal of Neuroscience, vol. 30, no. 50, pp. 17015–17022, Dec. 2010.

[9] J. T. Choi and A. J. Bastian, “Adaptation reveals independent control networks for human walking,” Nature Neuroscience, vol. 10, no. 8, pp. 1055–1062, Aug. 2007.

[10] R. J. Hamzey, E. M. Kirk, and E. V. L. Vasudevan, “Gait speed influences aftereffect size following locomotor adaptation, but only in certain environments,” Exp Brain Res, vol. 234, no. 6, pp. 1479–1490, Jun. 2016.

[11] T. Ogawa, N. Kawashima, T. Ogata, and K. Nakazawa, “Limited Transfer of Newly Acquired Movement Patterns across Walking and Running in Humans,” PLOS ONE, vol. 7, no. 9, p. e46349, Sep. 2012.

[12] T. Krasovsky and M. F. Levin, “Review: toward a better understanding of coordination in healthy and poststroke gait,” Neurorehabil Neural Repair, vol. 24, no. 3, pp. 213–224, Apr. 2010.

[13] R. Verma, K. N. Arya, P. Sharma, and R. K. Garg, “Understanding gait control in post-stroke: Implications for management,” Journal of Bodywork and Movement Therapies, vol. 16, no. 1, pp. 14–21, Jan. 2012.

[14] J. T. Choi, E. P. G. Vining, D. S. Reisman, and A. J. Bastian, “Walking flexibility after hemispherectomy: split-belt treadmill adaptation and feedback control,” Brain, vol. 132, no. 3, pp. 722–733, Mar. 2009.

[15] D. S. Reisman, H. McLean, J. Keller, K. A. Danks, and A. J. Bastian, “Repeated Split-Belt Treadmill Training Improves Poststroke Step Length Asymmetry,” Neurorehabilitation and Neural Repair, vol. 27, no. 5, pp. 460–468, Jun. 2013.

[16] J. M. Smoliga, E. J. Hegedus, and K. R. Ford, “Increased physiologic intensity during walking and running on a non-motorized, curved treadmill,” Physical Therapy in Sport, vol. 16, no. 3, pp. 262–267, Aug. 2015.

[17] A. M. Gonzalez et al., “Reliability of the Woodway Curve(TM) Non-Motorized Treadmill for Assessing Anaerobic Performance,” J Sports Sci Med, vol. 12, no. 1, pp. 104–108, 2013.

[18] C. J. Stevens, J. Hacene, D. V. Sculley, L. Taylor, R. Callister, and B. Dascombe, “The Reliability of Running Performance in a 5 km Time Trial on a Non-motorized Treadmill,” Int J Sports Med, vol. 36, no. 9, pp. 705–709, Aug. 2015.

[19] J.-C. Wang, W.-H. Sung, Y.-L. Chang, S.-H. Wu, and T.-Y. Chuang, “Speed and temporal-distance adaptations during non-motorized treadmill walking in stroke and non-disabled individuals,” Eur J Phys Rehabil Med, vol. 53, no. 6, pp. 863–869, Dec. 2017.

[20] R. T. Roemmich and A. J. Bastian, “Two ways to save a newly learned motor pattern,” J Neurophysiol, vol. 113, no. 10, pp. 3519–3530, Jun. 2015.

[21] G. Torres-Oviedo and A. J. Bastian, “Natural error patterns enable transfer of motor learning to novel contexts,” Journal of Neurophysiology, vol. 107, no. 1, pp. 346–356, Jan. 2012.

[22] R. S. Maeda, S. M. O’Connor, J. M. Donelan, and D. S. Marigold, “Foot placement relies on state estimation during visually guided walking., Control of Movement: Foot placement relies on state estimation during visually guided walking,” J Neurophysiol, vol. 117, 117, no. 2, 2, pp. 480, 480–491, Feb. 2017.

[23] J. S. Matthis, J. L. Yates, and M. M. Hayhoe, “Gaze and the Control of Foot Placement When Walking in Natural Terrain,” Current Biology, vol. 28, no. 8, pp. 1224–1233.e5, Apr. 2018.

[24] S. M. Morton and A. J. Bastian, “Cerebellar Contributions to Locomotor Adaptations during Splitbelt Treadmill Walking,” Journal of Neuroscience, vol. 26, no. 36, pp. 9107–9116, Sep. 2006.

[25] R. G. Ellis, K. C. Howard, and R. Kram, “The metabolic and mechanical costs of step time asymmetry in walking,” Proceedings of the Royal Society B: Biological Sciences, vol. 280, no. 1756, p. 20122784, Apr. 2013.

[26] N. Sanchez, S. Park, and J. M. Finley, “Evidence of Energetic Optimization during Adaptation Differs for Metabolic, Mechanical, and Perceptual Estimates of Energetic Cost,” Scientific Reports, vol. 7, no. 1, p. 7682, Aug. 2017.

[27] E. V. L. Vasudevan and A. J. Bastian, “Split-Belt Treadmill Adaptation Shows Different Functional Networks for Fast and Slow Human Walking,” Journal of Neurophysiology, vol. 103, no. 1, pp. 183–191, Nov. 2010.

