## Supplemental material: Secondary outcome for "Contributions of spatial and temporal control of step length symmetry in the transfer of locomotor adaptation from a motorized to a non-motorized split-belt treadmill"

Double support difference *(DS_diff)* is defined here as the difference between the fast and slow double support times. By calculating double support difference analytically, as with step length difference, the contribution of each individual component to the observed asymmetry can be assessed. Additionally, the relationship between the contributions of the foot placement, step time, and velocity can be compared between step length and double support difference to provide additional insight into how the locomotor pattern is being modulated during asymmetric walking conditions. Double support time, which will form the basis of our temporal difference measure, is defined as the difference between stance time and step time (Malone et al. 2012). To better understand how foot placement, step time, and belt velocity contribute to double support differences, an analytical derivation of double support difference was formulated for this experiment.

*Derivation of analytical double support difference.*

Here, we define double support time for the fast (slow) leading limb as the fast (slow) stance time minus the fast (slow) step time. The *leading limb* is defined here because both limbs are in contact with the treadmill surface during double support (by definition), however in asymmetrically constrained walking, double support times can differ significantly based on which limb is leading and the magnitude or ratio of belt speed differences. When calculating an analytical double support time, we wish to have individual components of foot placement difference, step time difference, and velocity difference to assess both their individual contribution to the measure as well as to compare them to those of the step length difference measure. To achieve this, we first define double support time as

$$DS = ST - t \tag{1}$$

where *ST* is the stance time and *t* is the step time for each limb during a step (Malone et al. 2012). Stance times, however, can be re-written as a function of limb excursion and belt velocity as follows

$$ST = \frac{LE}{v} \tag{2}$$

Limb excursion, the distance that a limb travels from heel strike to toe-off, can be written as the difference between the position of the foot at heel strike, or the forward foot placement (FFP), and the position of the foot at toe-off, or the rear toe-off position (RTP). However, since the accuracy of determining time and position of toe-off using the position of an ankle marker deteriorates with increasing plantar-flexion at toe-off, an analytical limb excursion is calculated which can then be subtracted from the forward foot placement to obtain the rear toe-off position. By combining equation (1) and equation (2) and rearranging terms, limb excursion is defined as

$$LE = v * (DS + t) \tag{3}$$

and now we can calculate the position of the foot at toe-off as the difference between the forward foot placement and the limb excursion

$$RTP = FFP - LE. \tag{4}$$

Now, inserting equations *2*, *3*, and *4* into equation 1, we can write our double support time as a function of forward foot placement, rear toe-off position, velocity, and step time as follows.

$$DS_f = \frac{FFP_f - RTP_f}{v_f} - \mathrm{t}_f \tag{5}$$

$$DS_s = \frac{FFP_s - RTP_s}{v_s} - \mathrm{t}_s\ . \tag{6}$$

This measure now describes the contribution of the foot placement, rear toe-off position, belt speed, and step time to the amount of time spent in double support for each limb on the respective belt. Therefore, we define the analytical double support difference as the difference between the fast and slow double support times

$$DS_{diff} = \frac{FFP_f - RTP_f}{v_f} - \frac{FFP_s - RTP_s}{v_s} + (t_s - t_f). \tag{7}$$

By rearranging terms, we can now write double support difference as a function of forward foot placement, rear toe-off position, step time, and belt speed differences

*(8)*

$$DS_{diff} = (t_s - t_f) + \left[\frac{v_f^{-1} + v_s^{-1}}{2} * (FFP_f - FFP_s)\right]$$

$$- \left[\frac{v_f^{-1} + v_s^{-1}}{2} * (RTP_f - RTP_s)\right] + \left[\frac{LE_f + LE_s}{2} * (v_f^{-1} - v_s^{-1})\right]$$

And simplifying, we arrive at a double support difference that is a function of differences in step time, foot placement, rear toe-off position, and the time to cover a meter.

$$DS_{diff} = \Delta t + \overline{v^{-1}} \cdot \Delta FFP + \overline{v^{-1}} \cdot \Delta RTP + \overline{LE} \cdot \Delta v^{-1} \quad (9)$$

Of note, two main differences evolved from the derivation of the double support difference analytical solution versus that of the step length difference analytical solution. First, there is a fourth term, the rear toe-off position difference, which is not present in the step length difference analytical solution. This should not be a concern because, first, both the foot placement and rear toe-off positions can be re-written as a difference in limb excursions, and second, because the double support time is merely the time between one foots heel contact and the other foots toe-off. Further, as can be seen in **figure S4**, the analytical double support difference perfectly predicts the instantaneous double support, despite that the rear toe position is calculated analytically. The last difference between the analytical derivation of step length and double support difference is that the velocities have been inverted to factor the double support difference into individual components. This may appear odd at first, however, it merely represents the speed component (and scaling factors) in terms of the amount of time taken to travel a meter (i.e. s/m) compared to the conventional distance covered in a second (i.e. m/s).

Repeated measures ANOVA revealed significant effects of time on double support ($F(7,63) = 57.25$, $p < 0.001$, **fig. S1a-S3a**). Group averaged double support differences were not significantly different from zero during baseline tied- slow, medium or fast walking on the motorized split-belt treadmill (all p-values > 0.05). **Figure S1a** shows gradual changes in double support difference (dark grey/black) during the first 100 and last 20 strides during Adaptation. During *early Adaptation*, double support difference showed a large initial change compared to baseline ($p < 0.001$, **fig. S1b,d**, black). By *late Adaptation*, double support difference was reduced back to baseline levels ($p = 0.66$, **fig. S1b,d**).

Like step length difference, the forward foot placement component for the double support difference showed an initial significant difference from baseline ($p < 0.001$) which reduced to near zero by around ten strides, then slowly increased to plateau by 100 strides. By *late Adaptation* the foot placement component was significantly greater than baseline ($p < 0.001$) and significantly greater than *early Adaptation* ($p < 0.001$, **fig. S1c,d**, red) despite *early Adaptation* being greater than baseline. The rear toe-off component was also significantly different from baseline ($p < 0.001$) during *early Adaptation*, made a large but non-significant increase by *late Adaptation* ($p = 0.08$), and remained greater than baseline ($p < 0.001$, **fig. S1c,d**, dark red). The step time component was not different from baseline during *early Adaptation* ($p = 0.24$) but increased to a plateau that was significantly greater than baseline ($p < 0.001$) and significantly different from *early Adaptation* ($p < 0.001$, **fig. S1c,d**, blue). The velocity component was significantly less than baseline ($p < 0.001$) during *early Adaptation*, made a significant decrease by *late Adaptation* ($p = 0.009$), and remained significantly different from baseline ($p < 0.001$, **fig. S1c,d**, magenta).

Consistent with step length difference, we found significant after-effects in double support difference during the transfer condition. Figure S2a shows a negative after-effect of double support difference which was significantly different from non-motorized treadmill baseline ($p < 0.001$) and returned to baseline symmetry by *late Transfer* ($p = 0.54$, **fig. S2a,b**). The foot placement component was no different from baseline at any time point ($p > 0.05$). The rear toe-off and step time components were both significantly different from baseline during *early Transfer* ($p < 0.001$), with the rear toe-off component less than baseline, and the step time greater than baseline. Both components returned to baseline by *late Transfer* (RTO, $p = 0.13$, ST, $p = 0.72$). The velocity component was significantly greater than baseline during *early Transfer* ($p = 0.04$) and made a significant decrease ($p = 0.003$) by *late Adaptation* that was significantly less than baseline ($p = 0.008$, **fig. S2c,d**).

Once again, robust after-effects in double support difference were apparent during *early Washout*. **Figure S3a** shows another robust negative after-effect of double support difference which was significantly different from the baseline slow treadmill condition ($p < 0.001$) which made a significant decrease ($p < 0.001$) back to baseline values ($p = 0.77$, **fib. S3b**). Similar again, to step length difference, the double support asymmetry was driven by initial differences in the foot placement component, which was significantly greater than baseline ($p < 0.005$) during *early Washout*. The foot placement component made a significant reduction ($p = 0.003$) back to baseline by *late Washout* which was not different from baseline ($p = 0.5$, **fig. S3c,d**). The rear toe-off and step time components were not different from baseline (RTO, $p = 0.47$, ST, $p = 0.08$), but quickly recovered a large asymmetry by about the tenth stride which took about 25-50 strides to recover the baseline symmetry. The rear toe-off component had a residual negative asymmetry which was significantly different from baseline ($p = 0.003$), while the step time component was not different from baseline ($p = 0.34$, **fig. S3c,d**). The velocity component was no different from baseline throughout the *Washout* period ($p > 0.05$).

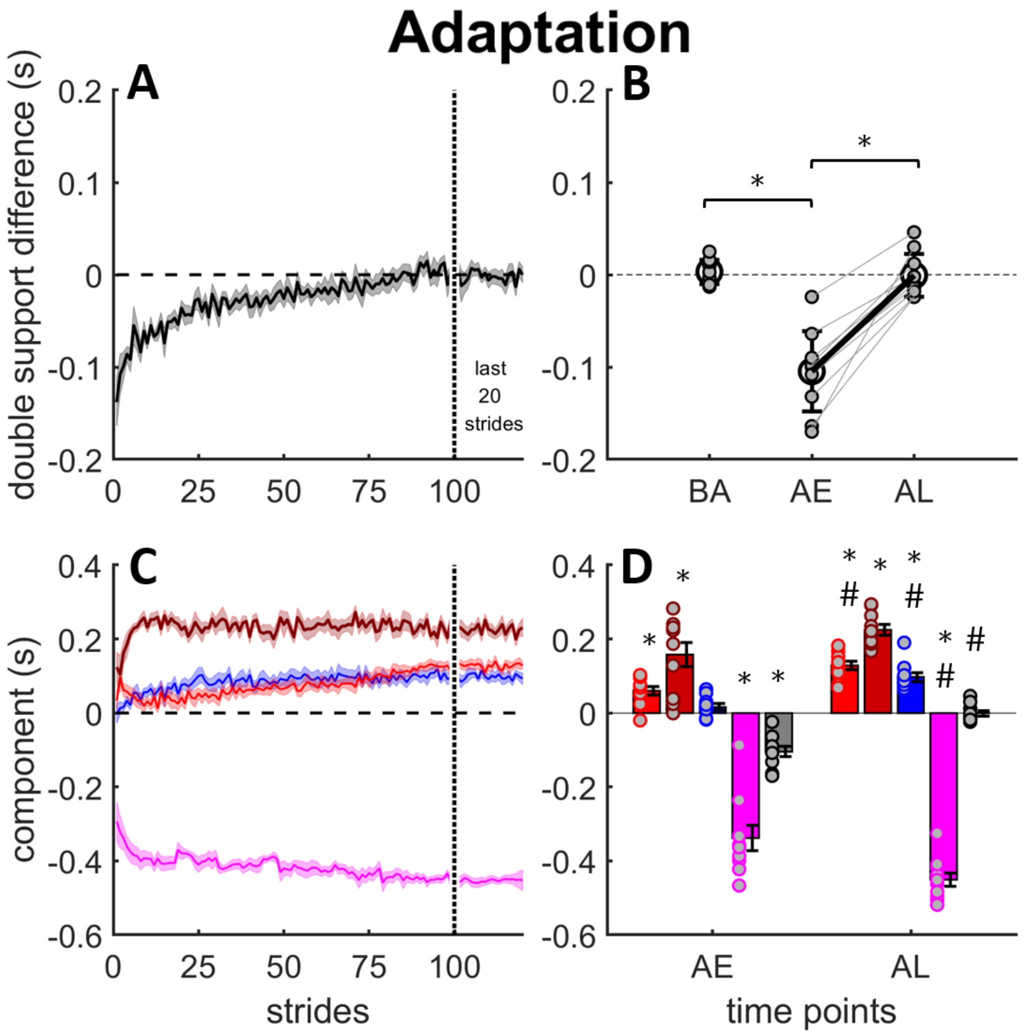


**Figure S1:** Adaptation; Progression of double support difference and contribution of spatial (foot placement and rear toe-off), temporal, and velocity asymmetries to motorized split-belt treadmill walking during adaptation. ***A.*** Group average double support difference during the first 100 and last 20 strides of the adaptation period. Shaded area represents plus/minus standard error. Negative values indicated the slow double support time is greater than the fast. ***B.*** Individual subject double support difference data (small gray filled circles) comparing baseline (BA) slow walking to early adapt (AE) and late adapt (AL). Light gray lines connect individual subject data points between AE and AL. Group means are represented by large black circles, error bars are standard deviations. Thick black line connecting AE and AL indicates group change. Horizontal bars with star indicate significant differences between time points ($p < 0.05$). ***C.*** Stride-by-stride changes in individual components for the first 100 and last 20 strides; foot placement (red), rear toe-off position (dark red), step time (blue), and velocity (magenta). Shaded areas are plus/minus standard error. ***D.*** Individual subject component data (gray filled colored circles) comparing AE and AL. Bars are group means, error bars are standard error. "*" indicates significant difference to baseline slow, "#" indicates late time point significantly different from early.

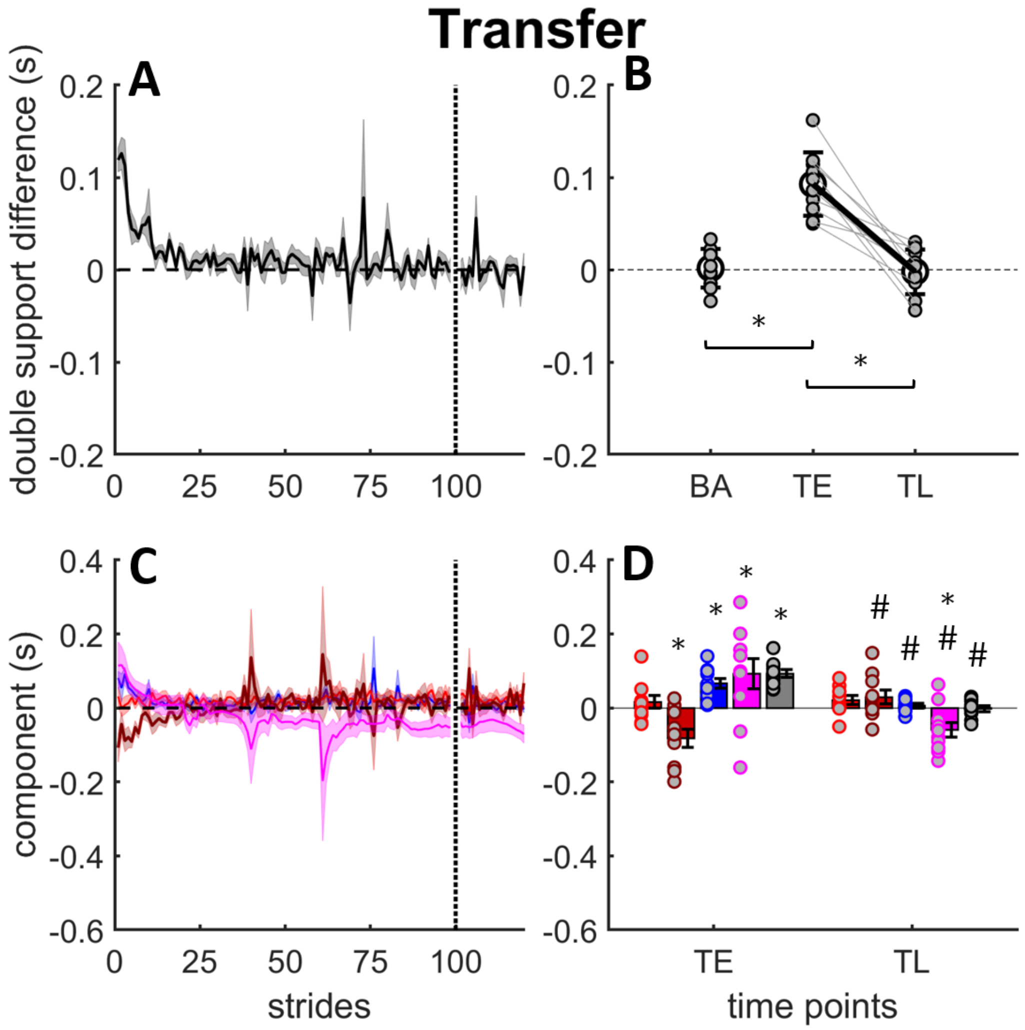


**Figure S2:** Transfer; Progression of double support difference and contribution of spatial (foot placement and rear toe-off), temporal, and velocity asymmetries to non-motorized split-belt treadmill walking during transfer. ***A.*** Group average step length difference during the first 100 and last 20 strides of the transfer period. Shaded area represents plus/minus standard error. Positive values indicated the fast double support time is greater than slow. ***B.*** Individual subject double support difference data (small gray filled circles) comparing baseline (BA) non-motorized treadmill walking to early transfer (TE) and late transfer (TL). Light gray lines connect individual subject data points between TE and TL. Group means are represented by large black circles, error bars are standard deviations. Thick black line connecting TE and TL indicates group change. Horizontal bars with star indicate significant differences between time points ($p < 0.05$). ***C.*** Stride-by-stride changes in individual components for the first 100 and last 20 strides; foot placement (red), rear toe-off position (dark red), step time (blue), and velocity (magenta). Shaded areas are plus/minus standard error. ***D.*** Individual subject component data (gray filled colored circles) comparing TE and TL. Bars are group means, error bars are standard error. "*" indicates significant difference to baseline non-motorized treadmill, "#" indicates late time point significantly different from early ($p < 0.05$).

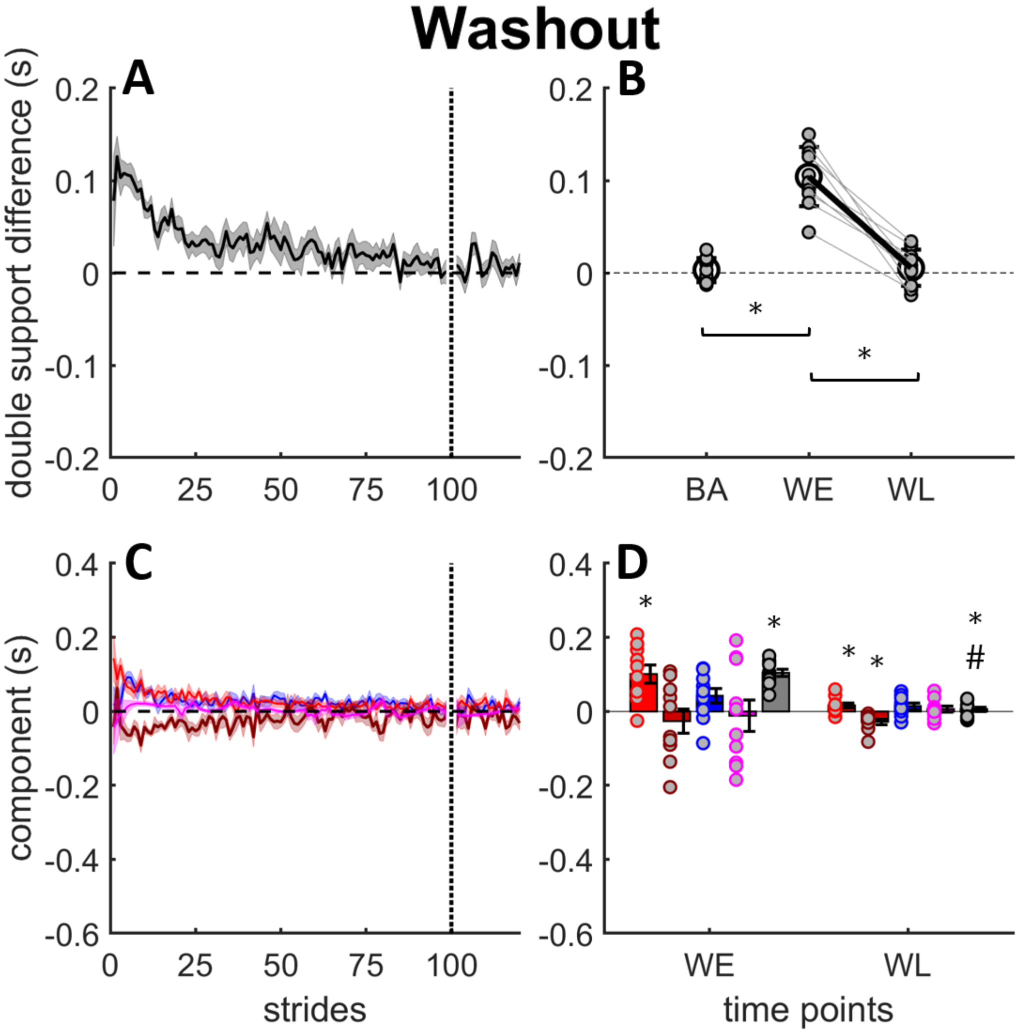


**Figure S3:** Washout; Progression of double support difference and contribution of spatial (foot placement and rear toe-off), temporal, and velocity asymmetries to motorized split-belt treadmill walking during adaptation. ***A.*** Group average double support difference during the first 100 and last 20 strides of the adaptation period. Shaded area represents plus/minus standard error. Positive values indicated the fast double support time is greater than the slow. ***B.*** Individual subject double support difference data (small gray filled circles) comparing baseline (BA) slow walking to early washout (WE) and late washout (WL). Light gray lines connect individual subject data points between WE and WL. Group means are represented by large black circles, error bars are standard deviations. Thick black line connecting WE and WL indicates group change. Horizontal bars with star indicate significant differences between time points ($p < 0.05$). ***C.*** Stride-by-stride changes in individual components for the first 100 and last 20 strides; foot placement (red), rear toe-off position (dark red), step time (blue), and velocity (magenta). Shaded areas are plus/minus standard error. ***D.*** Individual subject component data (gray filled colored circles) comparing WE and WL. Bars are group means, error bars are standard error. “*” indicates significant difference to baseline slow, “#” indicates late time point significantly different from early ($p < 0.05$).

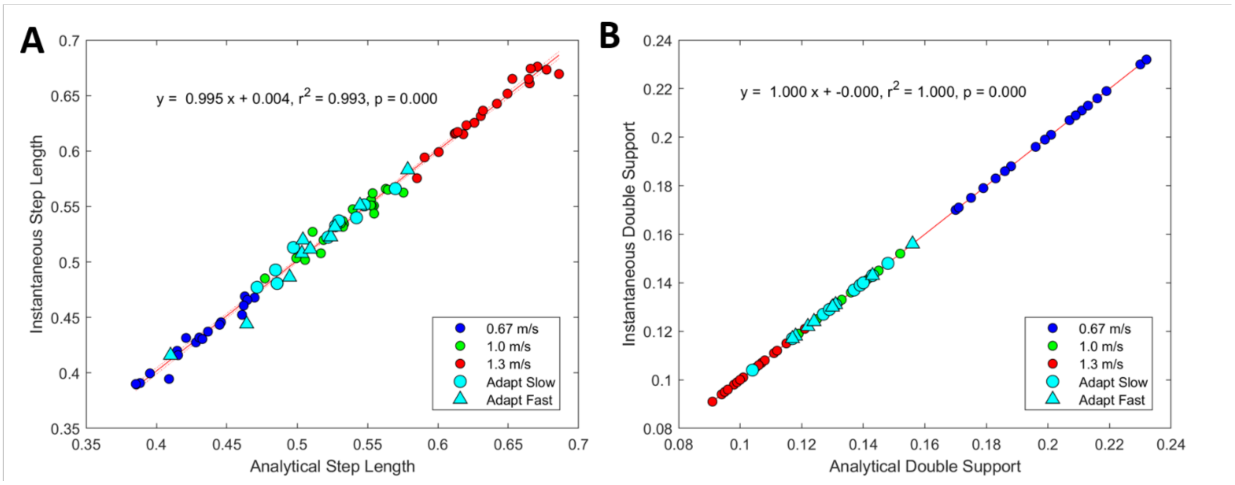


**Figure S4:** Comparison between analytical and instantaneous values for **A.** step length and **B.** double support time. Each data point is a the average step during steady state walking in slow (blue), medium (green), fast (red) and adapt (cyan) for each subject and for each limb, giving a total of 80 data points. The equation for the line of best-fit, $r^2$ and p-values are inset within each graph.
